# Absolute Quantification of Tissue Composition by Raman Spectroscopy

**DOI:** 10.64898/2026.10.25.761683

**Authors:** Erik E. Ersland, Dev R. Mehrotra, Henry Brisebois, Li-Hsin Han, Mark W. Grinstaff, Thomas Schaer, Mads S. Bergholt, Brian D. Snyder, Michael B. Albro

## Abstract

Raman spectroscopy captures the molecular fingerprints of living tissues without labels or tissue destruction. Translating these spectra into absolute measures of tissue biochemical composition remains a challenge owing to tissue-dependent optical properties, variations in sampling volume, and lack of normalization standards requisite for quantitative scaling. Here we introduce a scheme to derive the biochemical constituents of a tissue expressed as a percentage of the tissue wet weight, like biochemical assays, from Raman spectra measured optically. Nonlinear polynomial models of Raman spectra are trained on chemically defined phantoms that mimic the variability of the essential biochemical constituents of a tissue, accommodating spectral overlap and tissue-dependent optical effects. We validated this scheme using cartilage, a tissue fundamental to diarthrodial joint function. Cartilage is comprised of glycosaminoglycans, collagens and water that act synergistically to impart the rheological and tribological material properties essential to tissue function. Using biochemically perturbed live cartilage explants, phantom-calibrated Raman spectroscopy (PC-Raman) resolved constituent-specific changes and achieved quantitative agreement with independent biochemical assays (R² = 0.88-0.92; slopes ≈ 1.0; 95% confidence intervals ±2.7-3.2%pt). We demonstrate that PC-Raman is applicable across the hierarchy of tissue interrogation from microscopic imaging of tissue extracellular matrix composition to macroscopic longitudinal monitoring of drug effects *in-vitro* using live tissue constructs and *in-vivo* assessments of cartilage regeneration in an equine cartilage repair model. Converting Raman spectra measurements of the essential biochemical constituents of a tissue in absolute terms (tissue wet-weight-%) allows for the non-destructive, longitudinal quantification of tissue composition in disease (tissue degeneration) and in response to treatment (tissue repair).

## Introduction

Organ systems depend on the functional properties of the tissues that comprise them. Assessing tissue function in health and disease requires identifying standardized biomarkers that reflect tissue function and can be applied across a hierarchy of platforms: *in-vitro* studies on explanted tissue specimens, *in-vivo* preclinical small and large animal studies, and randomized clinical trials. Since tissue function is governed by the molecular building blocks and biochemical constituents of the tissue’s extracellular matrix (ECM), monitoring tissue ECM composition provides an objective biomarker for evaluating tissue function in health and disease. Advanced imaging modalities, such as compositional MRI (T1ρ, T2, T2*),^1;2;3^ ultrasound^4;5^, and optical coherence tomography^6^ portray tissue pathoanatomy but measurements of tissue composition using these modalities correlate only moderately with direct quantitative appraisals. Macroscopic tissue pathology grading schemes based on visualization of surface topography acquired using minimally invasive endoscopic systems are subjective^7^ and are poorly correlated with objective measures of tissue composition. Biochemical assays offer absolute measurements of ECM constituents but provide bulk averages for the sampled tissue and are destructive. Histopathological grading schemes are the gold standard for staging tissue pathology, but these evaluations are invasive and destructive, limiting repeat serial examinations over time and space. Histology staining depicts spatial changes in tissue composition and morphology for the sampled tissue but can be affected by processing and staining variability. Thus, there is need for a multiscale platform that can measure tissue ECM composition across the hierarchy of preclinical and clinical model systems from microscopic to endoscopic.

Raman spectroscopy is predicated on the inelastic scattering of photons by an incident monochromatic laser that induces wavelength changes that correspond to specific Raman active vibrational modes within a molecule^8^. The resulting Raman spectrum provides a quantitative, point-wise optical fingerprint of a tissue’s molecular building blocks (amides, sulfates, carboxylic acids, and hydroxyls), allowing non-destructive, label-free identification of the biochemical constituents comprising tissue ECM. Raman spectroscopy optical probes are compatible with hydrated tissues and can be integrated with a variety of imaging technologies including confocal microscopes and endoscopic systems enabling multi-scale *in-situ* tissue interrogation from microscopic to macroscopic across the hierarchy of model systems.

Converting Raman tissue spectra into absolute measures of biochemical composition presents a fundamental challenge. Raman analyses often use normalization routines to scale Raman spectral intensities, yielding relative spectral metrics that may not scale to absolute composition (**Fig 1A,B**). Normalization to an internal standard, reference band, or aggregate spectral intensity is performed to reduce measurement variability. However, this makes each signal dependent on a denominator that may change depending on the composition of the tissue sample. A decrease in one constituent’s signal artificially increases the normalized contribution of another without an actual change in its biochemical concentration (**Fig 1B**). Because of ‘spectral crosstalk’, the relative contribution of a constituent-specific Raman spectra profile to the overall tissue spectral profile may not scale linearly with the constituent’s mass fraction derived using defined physical parameter such as tissue volume or wet-weight as the denominator, independent of composition Spectral overlap presents another problem. While spectral deconvolution methods (including peak ratios, subpeak deconvolution, multivariate curve resolution^9^, N-Finder^10^, and reference-based multivariate regression^11^) can identify contributions associated with specific biochemical constituents, resolving their spectral contributions does not inherently relate to their biochemical concentration. Differences in Raman scattering efficiency, overlapping molecular signatures and tissue-dependent optical effects can produce nonlinear responses and additional crosstalk between constituent spectral profiles (**Fig. 1A,B**). To derive absolute measures of biochemical composition requires calibration of the tissue acquired Raman spectra against a standard of known composition.

**Figure 1:**
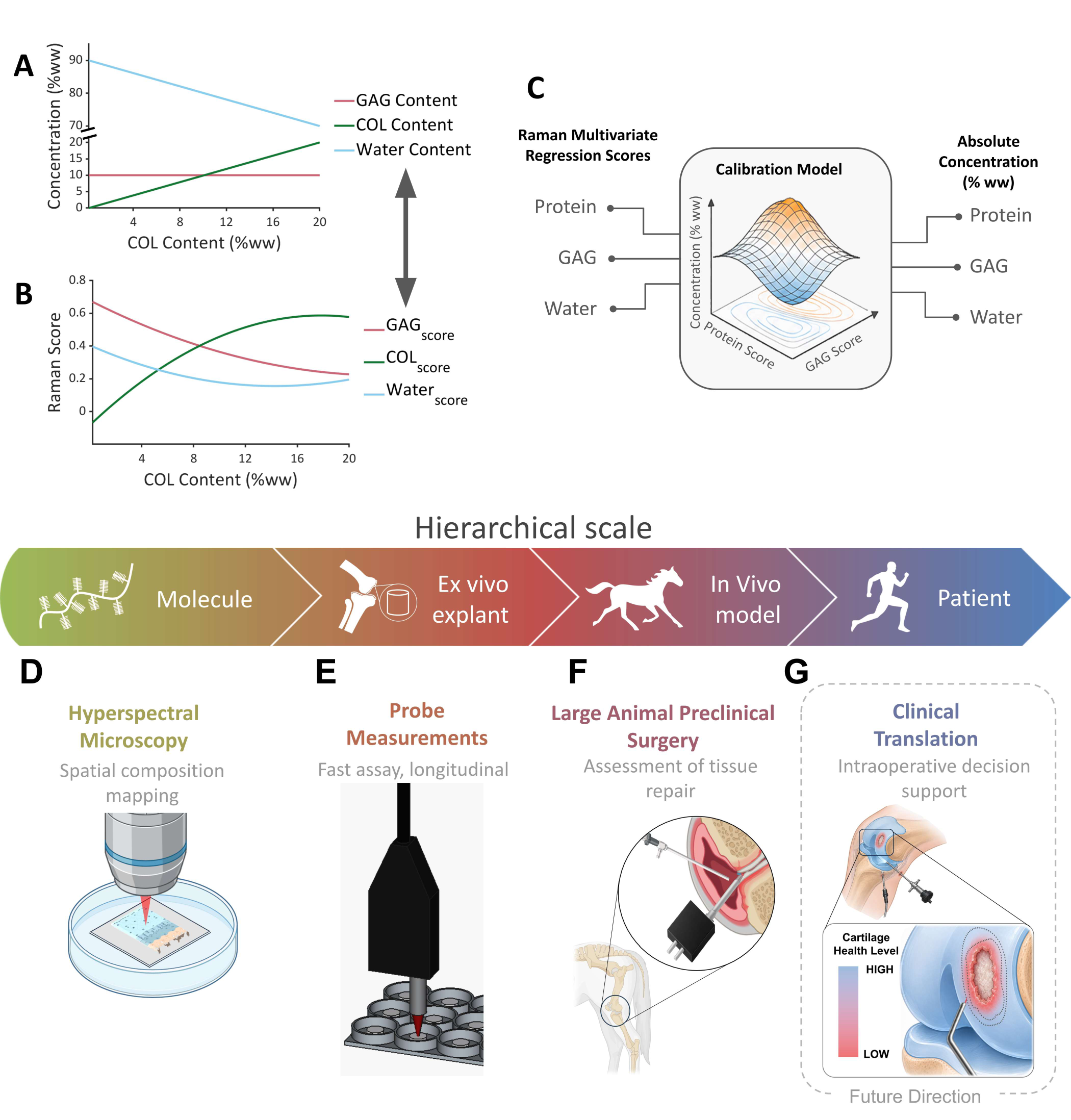
Calibration of Raman measurements for absolute quantification of tissue ECM. using cartilage, a tissue fundamental to diarthrodial joint function as an example. Cartilage is comprised of GAGs, collagens and water that act synergistically to impart the rheological and tribological material properties essential to tissue function. **(A,B)** Illustration of cross talk where changes in one component, such as COL content affects Raman scores of other components non-linearly. **(C)** Chemically defined phantoms (see Fig. 2 **A, D**) establish a nonlinear calibration that converts Raman scores into ECM constituent concentrations expressed as percentages of tissue wet weight. **(D–F)** Applications span **(D)** Raman microscopy for spatial compositional mapping, **(E)** probe measurements for serial assessment of live tissue **(F)** intraoperative measurements in an equine repair model, **(G)** Potential clinical applications are shown as a future direction.

***Our hypothesis is that tissue ECM composition and structure determine the functional material properties of a tissue, providing a biomarker accessible to Raman spectral analysis for monitoring tissue property changes in health, disease, and in response to regenerative therapies. The absolute content of specific ECM constituents important to tissue function can be calculated using tissue mimetic Phantoms to calibrate reference-based, multivariate least squares regression analysis of Raman spectral tissue profiles.*** Therefore, to convert Raman tissue spectra into absolute measures of biochemical composition, we developed and validated a framework predicated on external calibration of the acquired Raman tissue spectra using phantoms that mimic the variability of the essential biochemical constituents comprising a tissue in health and disease (**Fig. 2**). These chemically defined phantoms accommodate spectral overlap and tissue-dependent optical effects. A nonlinear polynomial model trained on the chemically defined tissue phantoms is used to convert the tissue acquired Raman spectra into the predominant biochemical constituents comprising the tissue ECM expressed as percentages of tissue wet weight (%ww). This scheme was validated using hyaline cartilage, a unique musculoskeletal tissue fundamental to diarthrodial joint function. Hyaline cartilage is a viscoelastic, biphasic, composite material comprised of a type-II collagen fibril network (5-20%ww) that provides tensile strength, complemented by a porous-permeable glycosaminoglycan (GAG) matrix (5-15%ww) of negatively charged, sulfated polysaccharides that regulate the transport of interstitial water (70-90%w/w)^12^. These ECM constituents synergistically impart the rheological and tribological material properties essential for cartilage articular function: support applied compressive and shear loads, while maintaining a nearly frictionless interface between articulating joint surfaces^13^.

**Figure 2:**
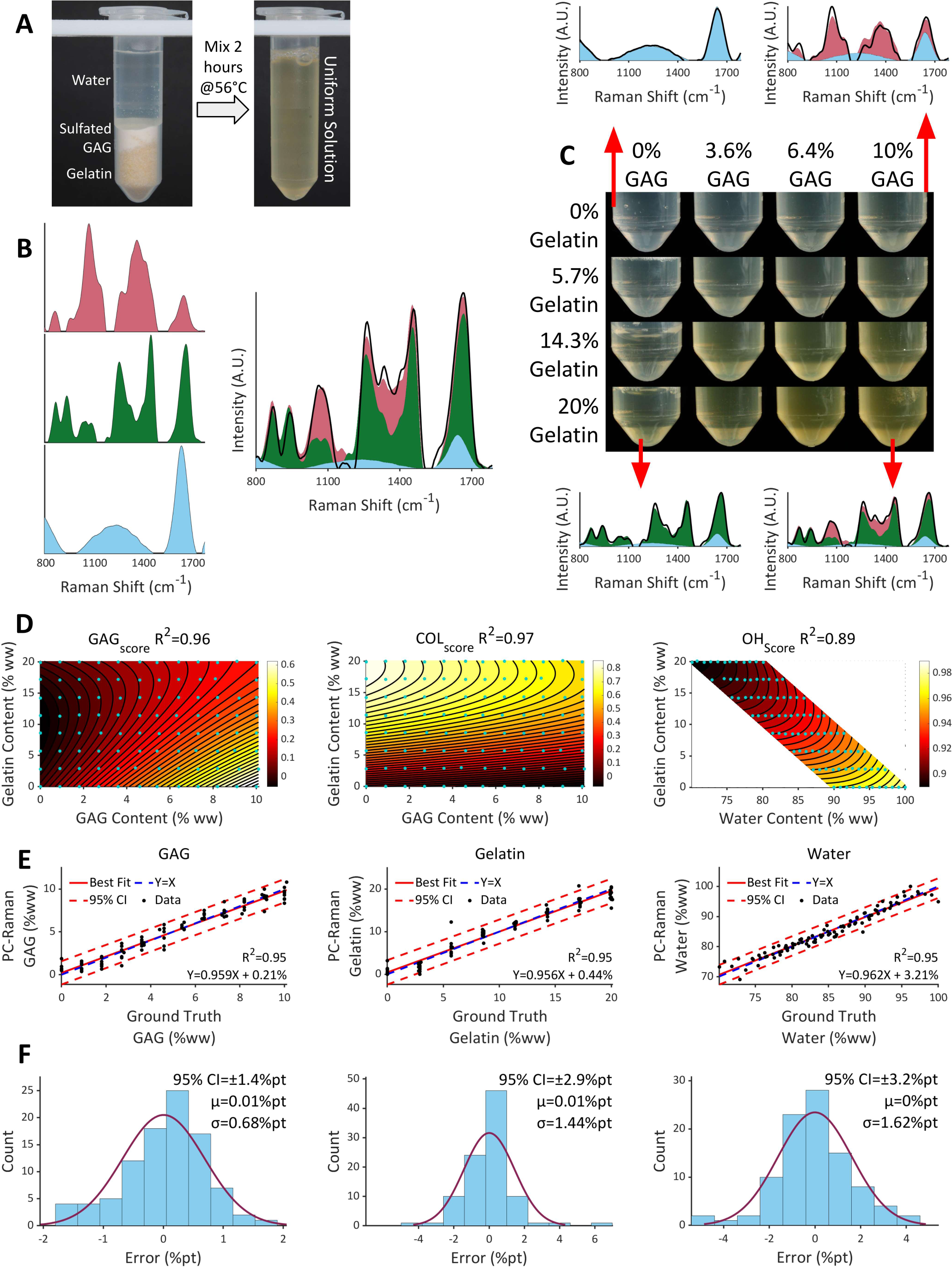
Tissue phantoms and calibration of Raman measurements. **(A)** Chemically defined phantoms prepared from chondroitin sulfate, gelatin and water. **(B)** Multivariate decomposition of fingerprint spectra (800–1800 cm⁻¹) yields constituent scores. **(C)** 2D array of phantoms span physiological composition ranges of 0–10% GAG and 0–20% protein. **(D)** Biomarker scores plotted as a function of ground-truth composition show nonlinearity and spectral crosstalk. **(E)** Nonlinear calibration converts these scores to constituent concentrations expressed as percentages of tissue wet weight. Leave-one-out cross-validation assesses predictions for GAG, protein and water. Parity plots compare predicted and ground-truth composition (R² = 0.95). **(F)** Error distributions are centered near zero, with reported 95% confidence intervals of ±1.4, ±2.9 and ±3.2 %pt for GAG, gelatin and water, respectively.

Osteoarthritis (OA) is a chronic, debilitating condition, characterized by the progressive degradation of hyaline cartilage manifest by matrix GAG depletion and derangement of the collagen network that result in increased tissue permeability and tissue swelling^14^, and decreased interstitial fluid load support and elastohydrodynamic lubrication that diminish the mechanical capacity of cartilage to withstand repeated compressive and shear loads during joint articulation^15^.

We prove the accuracy of “phantom calibrated” Raman spectroscopy (PC-Raman) to measure cartilage tissue ECM composition across the hierarchy of model systems, and conduct *in-situ* tissue interrogation over multiple spatial scales from microscopic to macroscopic using a confocal microscope to illustrate microscale compositional mapping and a probe to demonstrate non-destructive serial monitoring of live tissue explants *in-vitro* subjected to enzymatic degradation to emulate GAG depletion associated with OA and the therapeutic effects of insulin-like growth factor. *In-vivo* intraoperative assessment of cartilage spectra acquired with a 2-second integration time without sensitivity loss was accomplished using an arthroscopy compatible Raman optic needle probe that achieves high photon collection efficiency allowing non-destructive, real-time calculation of cartilage composition at discrete anatomic sites, including serial monitoring of neocartilage repair tissue in an equine stifle joint repair model (**Fig. 1D–F**). The derived GAG and water %ww were compared to standard biochemical assays for measuring GAG and water content in biological tissues. These experiments establish the capability of PC-Raman to perform across spatial scales and model systems.

## Results

### Tissue Specific Reference Phantoms for Calibrating Multi-variate Raman Spectral Analysis

To establish a quantitative relationship between Raman spectra and tissue composition, cartilage-specific reference phantoms were fabricated using solutions of chondroitin sulfate, the predominant GAG type in hyaline cartilage, and gelatin because it is spectrally similar to type-II collagen (**Supplement S2**), highly soluble and commercially available, and water (**Fig. 2A**). These ECM constituents are biomarkers that account for cartilage material properties. To mimic cartilage ECM in health and disease the compositions of the phantoms were varied parametrically over 12 uniformly distributed GAG concentrations from 0% to 10% and 8 uniformly distributed gelatin concentrations from 0% to 20%, with water comprising the remaining mass, yielding an array of 96 compositionally distinct reference phantoms (**Fig.2C**) (**Supplement S3**).

Raman spectra for the phantoms were acquired using a long-working-distance, contactless probe with the lens focused on the substance of the phantom, fiber-coupled to a near-infrared (NIR) diode laser and a spectrometer. Reference-based, multivariate-least-squares, non-negativity-constrained linear-regression analysis was performed on the fingerprint spectra (800-1800cm^-1^) for each phantom to derive regression coefficients (termed ‘scores’) GAG_score_, GEL_score_, H_2_O_score_ that reflect the relative contribution of each constituent (GAG, gelatin, and water) to the overall Raman fingerprint spectral profile for each phantom (**Fig. 2B**). Gaussian deconvolution of the high-wavenumber spectra (>2800 cm^-1^) yielded a water-content-associated OH_score_ and organic-content-associated CH_score_ ^16^(**Supplement S1**).

The GAG_score_, GEL_score_, and H_2_O_score_ varied nonlinearly with the known concentrations of the constituents contained in the phantoms (**Fig. 2D**). In particular, the relationship between GEL_score_ and gelatin concentration varied with GAG concentration. At high gelatin concentrations, the GEL_score_ became increasingly sensitive to GAG content and less sensitive to gelatin content. These coupled responses reveal why a constituent’s score cannot be a direct measure of a constituent’s concentration.

To calculate the concentration of a constituent from Raman scores requires deriving the relationship between the known concentration of the constituent in the phantom and the Raman scores (GAG_score_, GEL_score_, OH_score_) obtained by fitting a generalized second-order polynomial function with cross terms (*Score*_i_ × *Score*_j_) and 10 regression coefficients (*β*_ij_). This function represents the phantom calibration curve; the regression coefficients constitute the transformation matrix (**Equation 2**). Each constituent (GAG, gelatin, and water) has a unique and independent phantom calibration curve and transformation matrix. The best-fit model accounted for 95% of the variation in phantom composition (**Supplement S4**).

Uncertainty was analyzed using leave-one-out cross-validation (95 training phantoms iteratively tested against one held-out phantom). Notably, the Raman measurement error scaled as a constant percent of sample weight (**Fig. 2E**), instead of percent of the component being measured which is typical for a biochemical assay (**Supplement S5**). Error measures are reported as differences of weight percent or percentage points (%pt). The leave-one-out cross-validation (**Fig. 2F**) yielded normally distributed errors (error = Raman predicted chemical content - ground truth) with a mean of zero, allowing calculation of 95% confidence intervals (mean error ±2 standard deviations). The 95% confidence intervals for the concentration of each phantom constituent calculated using phantom calibration of reference-based multi-variate regression Raman scores are: ±1.4%pt for GAG, ±2.9%pt for gelatin, and ±3.2%pt for water. This indicates the lowest effect size that can be distinguished for cartilage ECM components using PC-Raman. A separate phantom calibration curve and transformation matrix was generated for each Raman hardware system: *in-vitro* tissue explant analysis using the contactless long-working-distance probe; microscopic analysis using confocal Raman microscope; *in-vivo* analysis using arthroscopy-compatible needle probe.

### Validation of Phantom Calibrated Raman Analysis in Native Tissue

The phantoms represent a simplified compositional model of native tissues that correct for spectral crosstalk and non-linearities inherent to conventional Raman analysis. They do not account for variation in ECM spatial distribution (e.g. compositional zonal gradients), microstructure, anisotropic structural organization, and other “minor” biochemical constituents that can confound Raman-derived measures of tissue composition. Since the phantoms use gelatin as a spectral surrogate for total collagen and does not reflect subtype, we denote the phantom calibrated protein spectra measured in cartilage as COL*. The efficacy of the 3-component phantom calibration in estimating cartilage ECM concentrations derived from reference based multivariate regression of Raman spectra acquired in real tissues was appraised in a parametric experiment conducted on live cartilage explants subjected to biochemical manipulation. Deep zone immature bovine cartilage explants were cultured for 0, 6, 10, or 14 days in medium supplemented with ±IL-1α, a catabolic cytokine that depletes cartilage GAG and collagen by shutting down the production of native cartilage matrix components and activating enzymes (aggrecanases and metalloproteinases) that break down both proteoglycans and collagen fibrils, leading to irreversible cartilage damage^17^ and ±dexamethasone (DEX), a synthetic corticosteroid that suppresses enzymatic ECM degradation^18^. After tissue explant exposures, Raman fingerprint spectra acquired using a long-working-distance contactless probe were subjected to reference based, multivariate regression to derive Raman biomarker scores (GAG_score_, COL_score_, H_2_O_score_) that portray the relative composition of the cartilage ECM. The concentration of each cartilage constituent (GAG, COL*, H_2_O) was calculated from the Raman scores using the phantom calibration curves and transformation matrices unique to each constituent of the cartilage ECM. These were compared to standard biochemical assays (wet/dry weight, DMMB colorimetric assay) and mechanical testing of cartilage compressive modulus.

Each treatment condition produced distinct changes in ECM composition and mechanical properties (**Fig. 3A-F**). The -DEX/-IL-1α condition produced COL* dilution and explant swelling driven by cell-secreted collagen-degrading enzymes in the absence of corticosteroid, while GAG content remained relatively stable^18^. The +DEX/−IL-1α condition served as the control condition with nearly stable composition and explant volume. -DEX/+IL-1α resulted in GAG depletion through IL-1α-mediated enzymatic degradation, accompanied by water replacement of the lost GAG. The +DEX/+IL-1α condition produced modest GAG loss due to partial inhibition of IL-1α activity by DEX. To generate other compositional variations, explants were subjected to delayed IL-1α exposure (±DEX) from day 14 to 21, eliciting GAG depletion after an initial swelling phase. These exposures generated a diverse range of physiologically relevant issue compositions: 0.3% to 10.3% GAG, 7.8% to 20.1% COL*, and 72.1% to 89.8% water (**Fig. 3G**). The minimal covariation between GAG and COL*(R^2^ = 0.24) allowed testing of constituent specificity. Enzymatic degradation of the tissue ECM adversely affected the material properties of the cartilage. Since compressive modulus depends on interstitial fluid load support, cartilage DMMB GAG and gravimetric water content jointly account for 94% of the variation in tissue modulus. Cartilage GAG and water content calculated using PC-Raman accounted for up to 93% of the variation in tissue modulus (**Fig.3H,I**).

**Figure 3:**
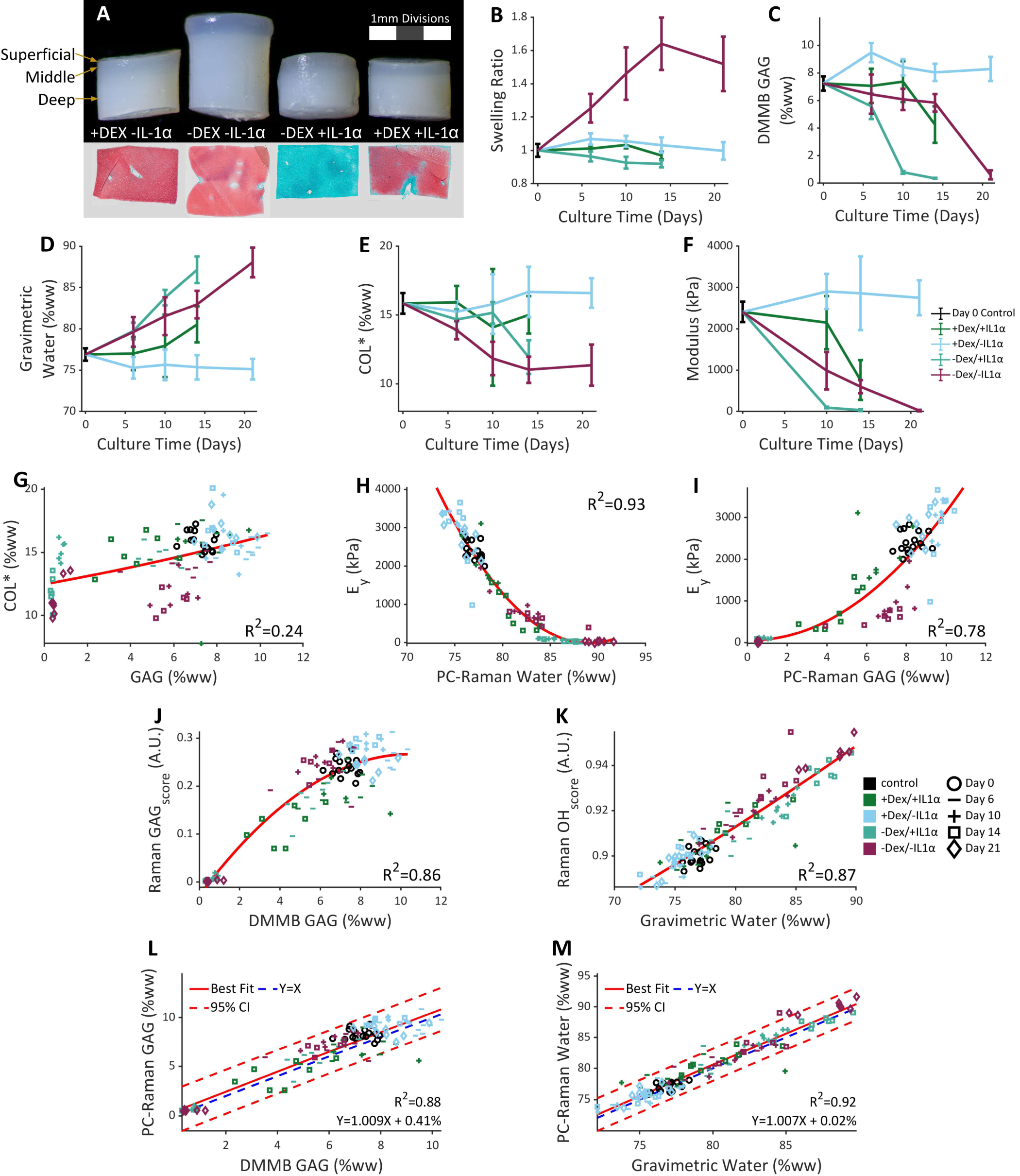
Validation of Raman quantification in live cartilage explants. **(A)** Explant morphology and corresponding histological sections stained with safranin O (red, GAG) and fast green (blue-green, collagen). **(B–F)** Time courses of biochemical composition and mechanical properties in immature bovine cartilage explants cultured under six conditions of IL-1α and dexamethasone (DEX) exposure. **(B)** Swelling ratio (weight normalized to control). **(C)** GAG content. **(D)** Water content**. (E)** COL* content**. (F)** Young’s modulus**. (G)** Validation samples span approximately 0–10% GAG, 8–20% protein and 72–90% water (n = 125 samples). **(H-I)** Composition-function relationship showing PC-Raman Water **(H)** and GAG **(I)** account for 93% and 78% of variation in modulus, respectively. **(J,K)** Conventional GAG and water scores compared with biochemical measurements (R² = 0.86 and 0.87, respectively). **(L,M)** Calibrated Raman measurements compared with biochemical measurements for GAG (R² = 0.88, slope = 1.01, intercept = 0.41 percentage points) and water (R² = 0.92, slope = 1.01, intercept = 0.02 percentage points).

As observed in our prior studies^19;20^ GAG_score_ and OH_score_ varied nonlinearly with direct measures of tissue content: DMMB measured GAG (R^2^ = 0.86) and water (R^2^ = 0.87) (**Fig. 3 J,K**) while the PC-Raman-derived tissue GAG and water content were linearly related: DMMB measured GAG (R^2^ = 0.88); water (R^2^ = 0.92) and nearly identical to the GAG and water content measured directly. The regression slopes were not different from unity (GAG: 1.01; water: 1.01), and the intercepts (0.41%pt and 0.02%pt were not different from zero; **Fig. 3L,M**). The calculated 95% confidence intervals were −1.7 to +2.7 %pt for GAG and −2.0 to +3.2 %pt for water, consistent with phantom calibration precision.

### PC-Raman Microscopy Quantitative Histology

Histology is the ‘gold standard’ for charactering cartilage pathology, but requires laborious sample processing, is subject to staining artifacts, and provides semi-quantitative measurement of cartilage ECM spatial composition. Phantom calibrated multivariate Raman spectral analysis enabled the generation of high-resolution quantitative microscopy images portraying the distribution of the major ECM constituents in tissue sections (**Fig. 4C, D**). Full-thickness live bovine cartilage explants (**Fig. 4A**) were serially exposed to IL-1α to produce progressive GAG depletion from the articular surface, or hyaluronidase to elicit severe GAG depletion. Explants were sectioned along the mid-sagittal plane. At each pixel, Raman fingerprint spectra were acquired over a 3 sec integration via the lens of the confocal microscope. Biomarker scores (GAG_score_, COL_score_, H_2_O_score_) derived from reference-based, multivariate regression analysis of the Raman spectra were used to calculate GAG, COL*, and H_2_O using the phantom calibration curves and transformation matrices unique to each constituent of the cartilage ECM. Untreated cartilage exhibited depth-dependent biochemical gradients, with lower GAG and higher water content in the superficial zone (SZ) and middle zone (MZ) and higher GAG and lower water in deep zone (DZ). Day 0 bovine cartilage sections displayed characteristic zonal GAG gradient: SZ = 1-4 %ww GAG; MZ = 4-6 %ww in (MZ); DZ = 6-10 %ww. IL-1α treatment induced progressive GAG depletion through the cartilage depth but COL* content remained relatively unchanged: 10%ww @ Day 0 and 11%ww @ Day 7 (not shown). Hyaluronidase produced immediate severe GAG loss throughout the entire specimen, with little effect on COL* content. Water content increased in regions with reduced GAG (**Fig. 4C**).

**Figure 4:**
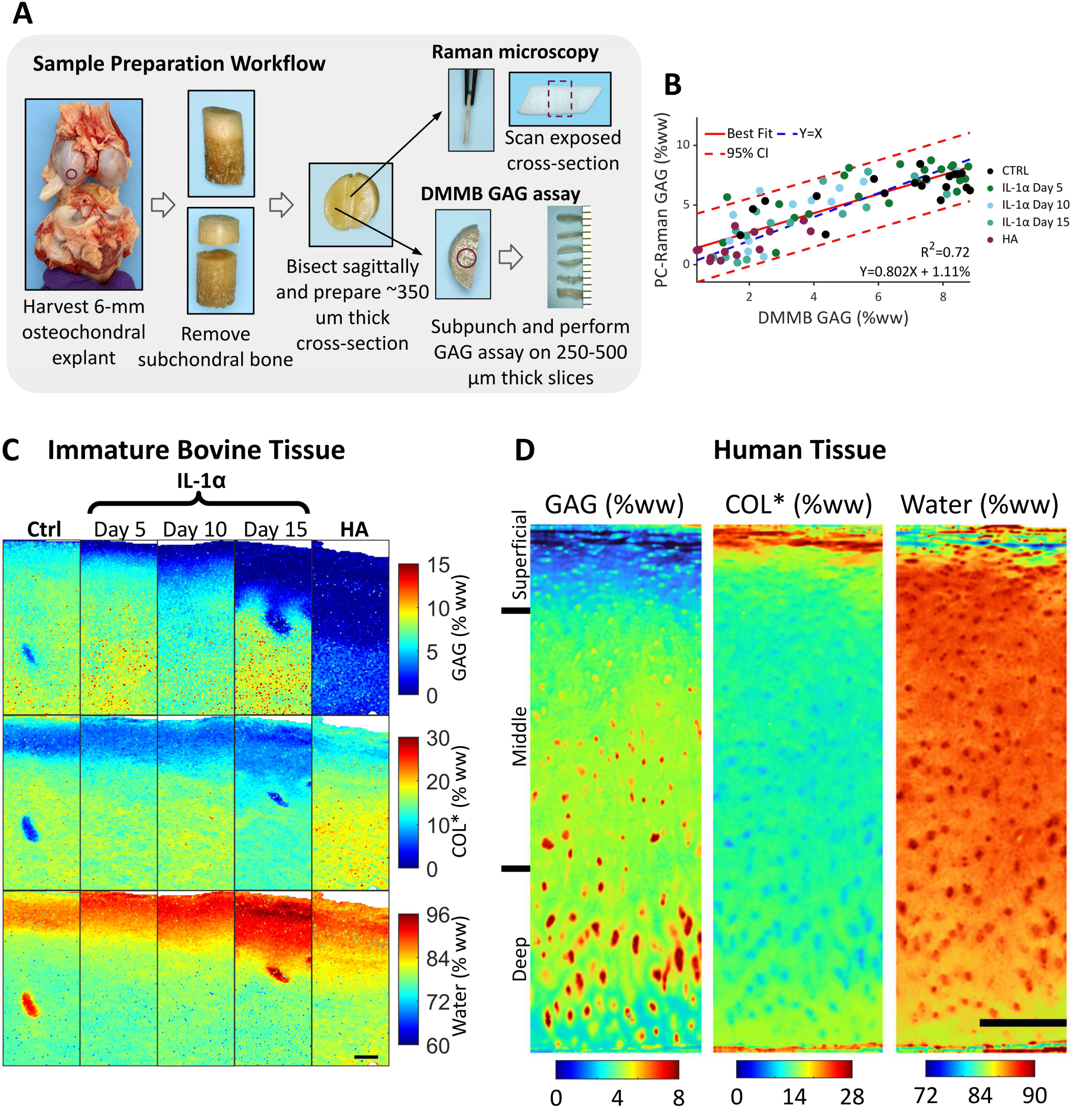
Spatial mapping of cartilage composition using PC-Raman quantitative microscopy. **(A)** Sample preparation workflow for paired Raman microscopy and DMMB GAG assay measurements from matched depth regions. **(B)** Raman-derived GAG content validated against biochemical GAG assay measures from paired specimens (R^2^ = 0.72). **(C)** PC-Raman-derived spatial distribution maps spanning from articular surface to 1.5 mm tissue depth. [Scale bar, 200 µm] for GAG, COL*, and water content in immature bovine cartilage cross-sections from control, IL-1α treated (5, 10, 15 days), and hyaluronidase treated explants. **(D)** Compositional maps of healthy human cartilage from a 19-year-old donor show depth-dependent composition gradients. [Scale bar, 300 µm].

PC-Raman-derived GAG content was compared to DMMB colorimetric GAG assay measurements from matched depth regions in adjacent tissue sections (**Fig. 4A**). PC-Raman derived GAG accounted for 72% of the variation in DMMB colorimetric GAG assay (R² = 0.72), with a 95% confidence interval from −2.8 to +3.2 percentage points. The best-fit linear regression between PC-Raman derived GAG content and DMMB colorimetric GAG assay content yielded a slope of 0.80 with an intercept of 1.1%pt (**Fig. 4B**).

Full-thickness human tibial plateau chondral explants harvested from 19-year-old human cadaver knee were fixed overnight in formalin, bisected, embedded in OCT and cryosectioned to 100µm slices, mounted on steel plates, and submerged in PBS for Raman confocal microscopic imaging. Consistent with the ECM compositional gradients that characterize healthy mature cartilage spatial distribution maps of GAG, COL*, and H_2_O content demonstrate quantitatively low-GAG and high-water content at the SZ, increasing GAG content with depth and relatively uniform COL* content through the MZ (**Fig. 4D**). These label-free PC-Raman generated high-resolution quantitative microscopy images depict the spatial distribution of the major ECM constituents in tissue sections like standard histology without the need for special preparation or histological stains, thereby preserving the tissue for other types of analyses on the same tissue sections.

### PC-Raman Quantification of Compositional Changes Cartilage Explants *In-vitro*

Raman provides a pragmatic preclinical tool for screening the efficacy of novel therapies *in-vitro.* Live immature bovine femoral cartilage explants were treated with IL-1α ± insulin-like growth factor 1 (IGF-1), a candidate therapeutic agent that promotes cartilage preservation by stimulating chondrocyte proliferation, enhancing ECM production, and inhibiting chondrocyte apoptosis^21^. Using Raman-compatible sterile tissue culture chambers capable of maintaining a sterile contact-free workflow requisite for long-term cartilage explant viability, longitudinal, repeated-measure of Raman fingerprint spectra were acquired via a long-working-distance contactless probe through the well-plate window (**Fig. 5A, B**). The spectra were subjected to reference based, multivariate regression to derive Raman biomarker scores (GAG_score_, COL_score_, H_2_O_score_) that portray the relative composition of bulk tissue cartilage ECM and site-specific ECM heterogeneities. The concentration of each cartilage constituent (GAG, COL*, H_2_O) was calculated from the Raman scores using gelatin phantom calibration curves and transformation matrices unique to each cartilage ECM constituent. Explants subjected to IL-1α over 10 days of culture displayed ∼61% GAG depletion in association with increased water content; COL* content remained stable over 10 days, consistent with prior work showing that GAG depletion precedes collagen loss^22^(**Fig. 5E**). On day 10, control explants displayed significantly greater GAG content than IL-1α-treated explants (P < 0.01). IGF-1 treatment partly mitigated (∼40%) the IL-1α-induced GAG loss and concomitant water accumulation (**Fig. 5F**). For explants cultured under identical conditions, variation in PC-Raman-derived GAG and water content accounted for 82% and 64% of the variation in DMMB measured GAG (**Fig. 5C**) and gravimetric water content (**Fig. 5D**), respectively (95% confidence intervals: −2.0 to +2.5 %pt for GAG; −3.5 to +4.0 %pt for water).

**Figure 5:**
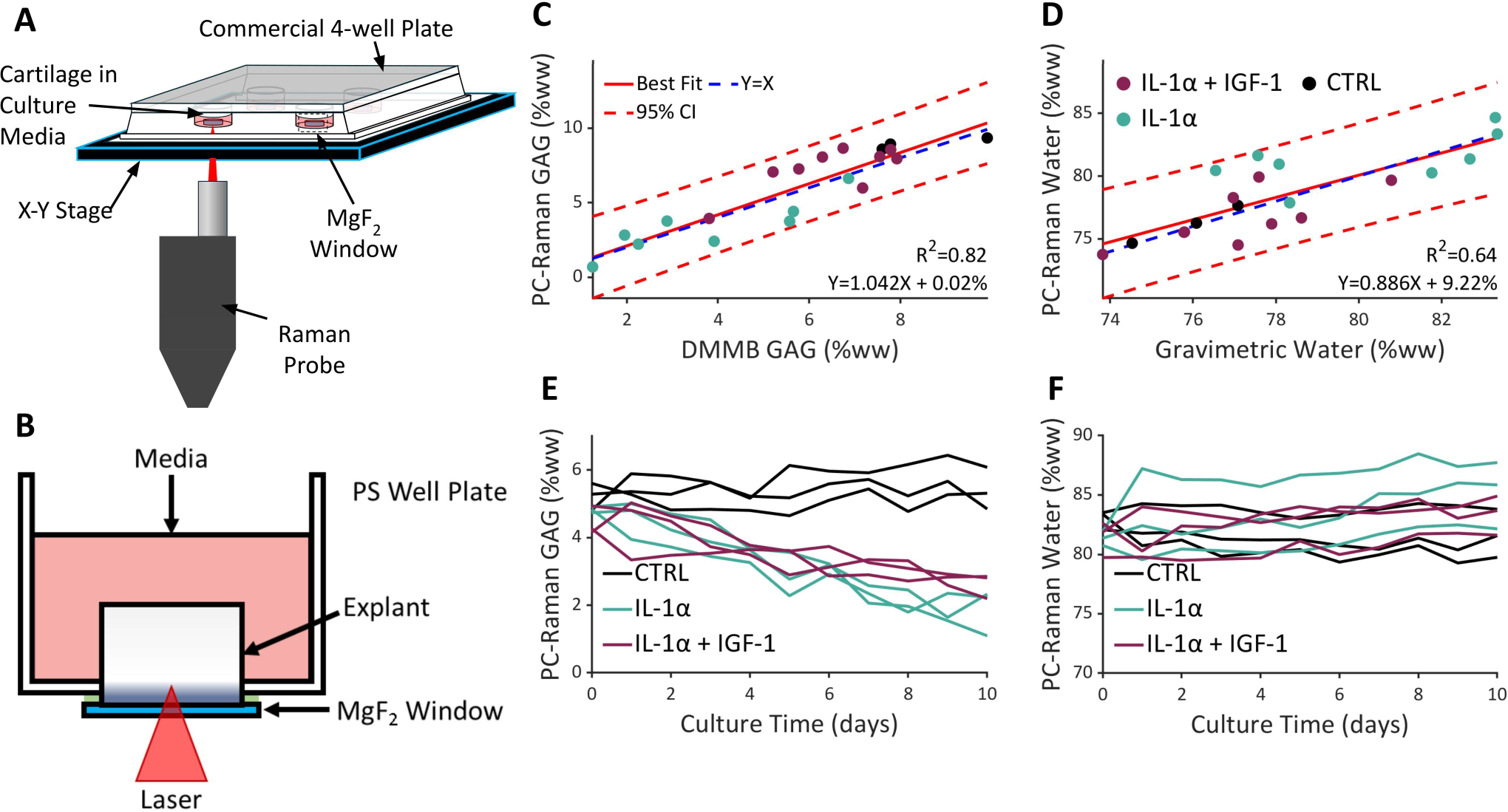
Serial Raman quantification of drug-induced composition changes in live cartilage. **(A)** Raman spectra were acquired using a contactless long-working-distance probe imaging through a custom well plate. **(B)** Section view of customized culture well with Raman-transparent MgF2 windows enabled non-destructive daily measurements through the well bottom. Samples were imaged superficial side down. **(C-D)** Endpoint validation: PC-Raman-derived measures versus biochemical assays from parallel explants showing **(C)** GAG (R^2^ = 0.82) and **(D)** water (R^2^ = 0.64) content. **(E-F)** Serial Raman measurements over 10 days showing **(E)** GAG and **(F)** water content changes in explants cultured under control, IL-1α, and IL-1α + IGF-1 treatment.

### PC-Raman Quantification of Compositional Changes Neocartilage *In-vivo,* Equine Stifle Joint model

PC-Raman achieved *in-vivo*, intraoperative monitoring of chondroregeneration in a preclinical large animal model analyzing the formation and maturation of neocartilage formed in an osteochondral defect. Raman spectra were acquired using an arthroscopy-compatible needle probe with a distal ∅2 mm sapphire ball lens (**Fig. 6A**) held normal to the cartilage surface with minimal contact force (**Fig.6D**). Two 15-mm-diameter chondral defects were created at proximal and distal poles of the lateral trochlea facet and treated by cell seeded collagen microfilament scaffolds (**Fig. 6C**). Tissue spectra were analyzed with a 2 sec integration time at multiple sites within each defect and the surrounding native cartilage, plotted in a polar coordinate reference system (**Fig. 6B**). Serial *in vivo* assessments were performed at baseline (before defect creation) and at 3, 6 and 12 months after-repair (postmortem) (**Fig. 6E**). Visual inspection and Raman revealed neocartilage filling the volume of the defect. The concentration of each cartilage constituent (GAG, COL*, H_2_O) was calculated from the Raman scores using gelatin phantom calibration curves and transformation matrices unique to each cartilage ECM constituent. The presence of the collagen scaffold contributed to elevated COL* content in the defect over time, matching or exceeding native tissue COL* levels. However, the GAG content of the neocartilage remained substantially below that of the excised cartilage it is replacing or the surrounding native cartilage. Thus, serial Raman interrogation of the neocartilage repair tissue indicated that retention of the collagen microfilament scaffold contributed significantly to the fibrocartilaginous tissue repair, and that the seeded chondrocytes produced very little GAG requisite for interstitial fluid load support and elastohydrodynamic lubrication^15^(**Fig. 6F**). While the repair tissue filling the volume of the defect appeared to be robust, the neocartilage did not reconstitute the ECM constituents essential for cartilage articular function to withstand repeated compressive, and shear loads during joint articulation. A real time software interface was then developed which can display phantom calibrated readings in as little as 0.4 seconds (**Supplement S6**).

**Figure 6:**
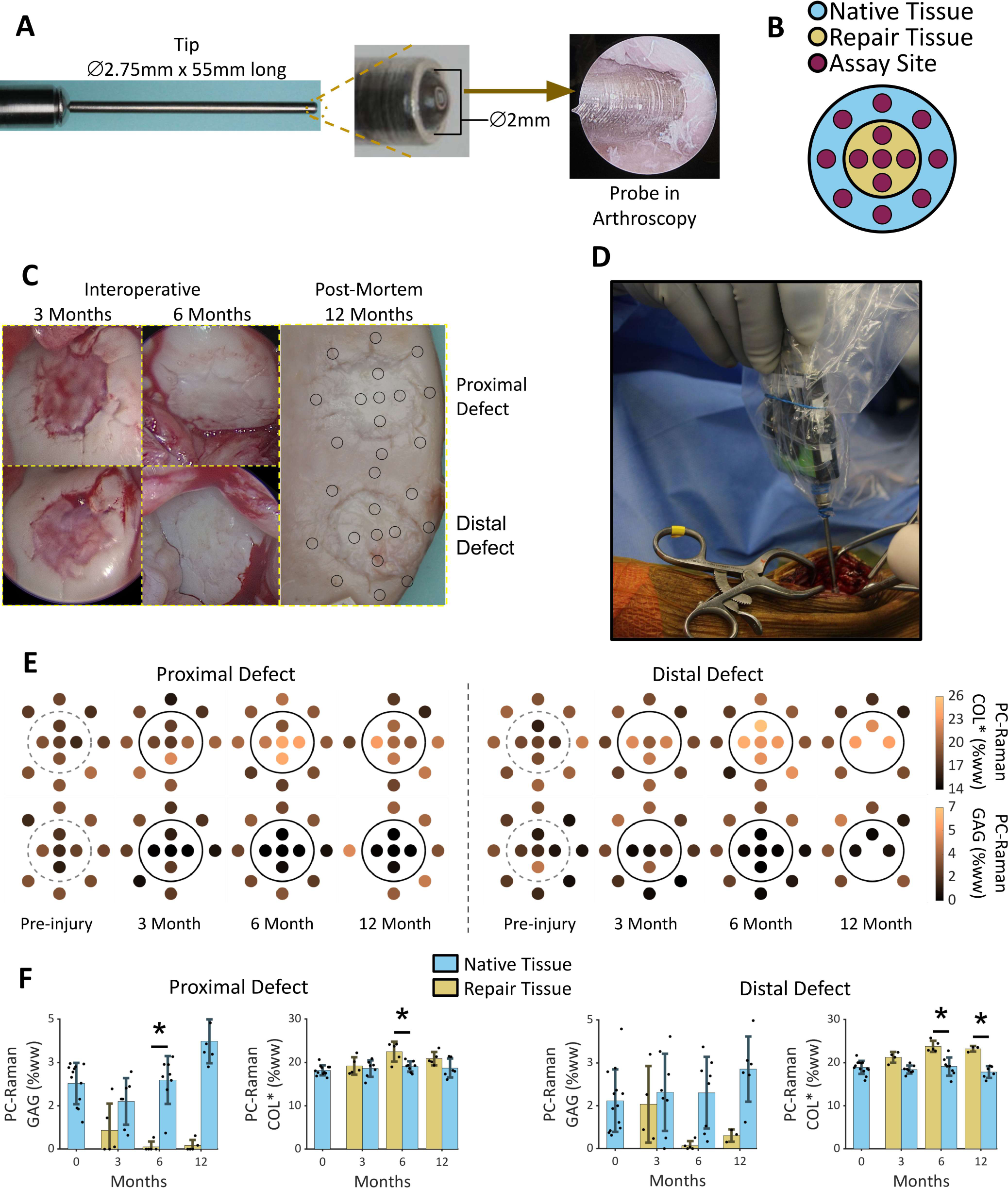
Longitudinal Raman assessment of cartilage repair in an equine model. **(A)** Raman spectra were acquired using an arthroscopy-compatible needle probe with a ⌀ 2.75mm bore and ⌀2mm lens. **(B)** Polar coordinate reference system for Raman measurement locations within repair tissue and surrounding native cartilage. **(C)** Photographs of cartilage repair sites at 3 and 6 months *in vivo* and at 12 months postmortem, with measurement locations marked on the final image. **(D)** Intraoperative probe positioning for Raman measurement of native tissue sites. **(E)** PC-Raman-derived GAG and COL* content distribution at baseline and post-repair time points (3, 6, 12 months). Large circles mark defect boundaries. **(F)** Raman-derived GAG and COL* content over time comparing repair sites versus surrounding native cartilage. *p<0.05 indicates significant difference. Error bars indicate standard deviations.

## Discussion

Raman spectroscopy enables label-free, non-destructive tissue analysis with high molecular specificity, yet conventional methods generally yield semi-quantitative measurements that are difficult to compare across laboratories or instruments, limiting their clinical and research utility. Phantom calibration places Raman measurements on an absolute biochemical scale, enabling absolute measures of biochemical constituent concentrations in tissue directly comparable to gold-standard biochemical assays. This approach provides a non-destructive, label-free, quantitative biochemical assessment platform applicable across spatial scale, from microscopic benchtop to macroscopic *in-vivo* surgery, and across the hierarchy of tissue model platforms, from *in-vitro* studies on explanted tissue specimens to *in-vivo* preclinical animal studies, and randomized clinical trials, advancing the translational potential for Raman spectroscopic assessment of tissue biomarkers that are critical to tissue function.

Spectral crosstalk between biochemical constituents is a confounding factor that limits the quantitative utility of conventional spectral unmixing algorithms. Reference-based, multivariate least squares, non-negativity-constrained linear regression of Raman fingerprint spectra constrains the derived regression coefficients (scores) representing the relative contribution of the predominant constituents of the tissue ECM to sum to unity. Consequently, an apparent increase in one constituent necessarily produces a corresponding decrease in others, complicating quantitative interpretation of tissue ECM composition. Just as calibration curves enable absolute quantification in standard biochemical assays, PC-Raman employs chemically defined reference phantoms as calibration standards to convert the reference based multivariate Raman spectral tissue profiles into absolute concentrations of ECM constituents expressed as percent tissue wet weight. This transformation was achieved by fitting a generalized second-order polynomial model incorporating constituent-specific terms and cross terms to account for nonlinear relationships and interactions among spectral contributions. The polynomial represents the calibration curve, and the regression coefficients constitute the transformation matrix for the phantom (Equation 2). The best-fit model accounted for 95% of the variation in the 3-component phantom and specified estimates of accuracy and precision across the calibration range.

The live cartilage explant culture model offered physiologically relevant validation of PC-Raman in complex tissues. Systematic chemical perturbations through cytokine and corticosteroid manipulation elicited distinct orthogonal compositional changes: IL-1α selectively decreased GAG, and in the absence of corticosteroid caused tissue swelling and decreases in COL*. A test of the specificity and sensitivity of PC-Raman measurements was the minimal covariation between GAG and COL* (R^2^ = 0.24), marginalizing the risk that a metric associated with one constituent appears to be predictive of others when tissue constituents change together. PC-Raman analysis quantified GAG and water across perturbed compositional states in agreement with independent biochemical assays (R^2^ = 0.88-0.92), supporting the hypothesis that the content of specific ECM constituents important to tissue function can be calculated using tissue mimetic phantoms to calibrate reference-based, multivariate least squares regression of Raman spectral tissue profiles.

PC-Raman was successfully applied across experimental and clinically relevant settings. For microscopy, PC-Raman enabled generation of micron-resolution quantitative tissue histology without fixation or staining, avoiding processing artifacts and staining variability. For *ex-vivo* studies, PC-Raman offered technical advantages over conventional biochemical assays that require sample destruction and laborious processing of tissue specimens (e.g., enzymatic digestion and colorimetric analysis). PC-Raman was non-destructive, with minimal preparation of samples for bulk measurements enabling: 1) serial monitoring of live tissue explants in response to mechanical, chemical, and pharmacological perturbations, and 2) high-throughput composition screening of large sample sets with reduced labor and materials costs. For *in vivo* preclinical animal studies that monitor therapeutic response to chondroregenerative procedures, PC-Raman can deliver real time, minimally invasive “optical biopsy” of effected tissues after simulated injury, tracking longitudinal changes in tissue composition that account for tissue function. The equine study demonstrates that PC-Raman can be used in clinical trials where cartilage is accessed arthroscopically to quantify therapeutic efficacy of chondroregenerative or chondroprotective strategies where the therapeutic goal is to rejuvenate injured or diseased cartilage to a healthy compositional state.

Standard-curve calibration of Raman spectra has precedent in other disciplines, including mineralogy^23;24;16^, pharmaceutics^25;26;27^, and ASTM standards for petroleum products^28^. Previous work has also extended calibration-based Raman analysis to biological systems. In particular, the approach termed *qRamanomics*^29,30^ demonstrated submicron-resolution, three-dimensional Raman microscopy of highly cellular organoids using complex phantom fabrication procedures to quantify multiple biochemical constituents. The present study builds upon these foundational advances by introducing a comparatively simple tissue-mimetic phantom fabrication strategy, establishing quantitative agreement with independent biochemical assays in live tissue, and demonstrating applicability across microscopic and macroscopic measurement platforms. These advances extend the utility of calibration-based Raman spectroscopy beyond specialized microscopy applications toward broader tissue characterization and translational research.

Chemically defined biochemical phantoms also provide a foundation for improving the reproducibility and standardization of quantitative Raman spectroscopy. By establishing known, reference compositions, phantoms can support: 1) evaluation and optimization of spectral processing algorithms, including autofluorescence baseline subtraction and multivariate unmixing routines; 2) validation of hardware performance for measurement accuracy, and 3) standardization across different laboratories and clinical sites using calibrated Raman measurements to generate quantitatively comparable results across hardware configurations (e.g., spectrometer resolution or detector sensitivity), addressing a fundamental barrier to clinical translation. Although cross-instrument standardization was not directly evaluated in the present study, the use of chemically defined reference materials provides a potential route toward harmonizing quantitative measurements across different hardware configurations.

Opportunities remain to extend the scope and generalizability of the PC-Raman framework. Future work will incorporate additional biochemical constituents relevant to other tissues, including mineralized components of subchondral bone, elastin-rich matrices for tendon and lung, and lipids for liver^30;31;32^. Furthermore, although the present study employed reference-based multivariate spectral regression, the general principle of phantom-derived calibration is not inherently restricted to this analytical approach. Other spectral analysis methods that generate composition-sensitive outputs, including peak intensity ratios, principal component-based models, and machine learning algorithms, could potentially be calibrated against chemically defined phantoms to recover absolute biochemical concentrations. Collectively, these developments could broaden the application of PC-Raman as a non-destructive platform for quantitative biochemical assessment of tissue composition in disease, regeneration, and therapeutic intervention

## Methods

### Raman instrumentation

#### *In-vitro/Ex-vivo* Explant analysis

Raman spectra were acquired using a contactless long-working-distance probe (In Photonics model RPB785) fiber-coupled to a NIR diode laser (excitation wavelength = 785 nm; IPS model 10785MU0500MS-USB, delivering 85 mW to tissue) and a spectrometer (Ocean Optics model QEPRO 01497; 0 to 4050 cm^-1^;1044 pixels). The probe exhibits a laser spot diameter of ∼380 um with a 7.5 mm working focal distance. All spot sizes were measured with a CCD in the focal plane.

#### Confocal microscopy analysis

Raman spectra were acquired on a custom confocal Raman microscope^33^ equipped with a 785 nm laser (120 mW; Toptica IBeam), transmitted through a 60× water immersion objective (Nikon), motorized stage (Zaber), and fiber-coupled spectrometer (Ibsen model Eagle DR-316B-LDC-DD, 100-3400 cm^-1^, 2000 pixels).

#### *In-vivo* analysis

Raman spectra were acquired using an arthroscopy-compatible needle probe (In Photonics custom model RPC785-PP-C)^34;35^ with a threaded needle tip (∅2.75 mm OD) containing a distal ∅2 mm sapphire ball lens (**Fig. 6A**). The probe generates ∼550 µm spot size with exponential decay depth sensitivity greatest at the surface, decreasing by a factor of 2 for each 470 µm of cartilage depth. The probe is fiber-coupled to an NIR diode laser (785 nm, IPS model 10785MU0500MS-USB delivering 150 mW to the tissue) and high-performance spectrometer (Ibsen model Eagle DR-316B-LDC-DD, 0-3700 cm^-1^, 2000 pixels).

### Multivariate Raman spectral analysis

Raman spectra were preprocessed according to the NIST SRM 2241 certificate specifications, including wavenumber calibration against polystyrene (ASTM E1840-96), probe background subtraction, and absolute intensity correction. The spectral fingerprint range (800-1800 cm^-1^) and high-wavenumber range (>2800 cm^-1^) were cropped and separately preprocessed through autofluorescence removal (fingerprint: 4^th^-order polynomial; high-wavenumber: exponential decay). The fingerprint region was subjected to reference-based, multivariate least squares, non-negativity-constrained linear regression^19,20^:

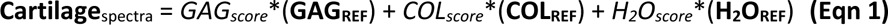

where the derived regression coefficients (*GAG_score_, COL_score_, H_2_O_score_*) represent the relative contribution of the predominant constituents of cartilage ECM (GAG, COL, H_2_O) to the composite cartilage Raman spectra (**Fig.2B**), calculated using the spectra of purified reference chemicals for each ECM constituent: **GAG_REF_**=chondroitin sulfate (bovine cartilage; Sigma[C6737]); **COL_REF_**=type-II collagen (chicken sternal cartilage; Sigma[C9301]); **H_2_O_REF_**=deionized water. For each constituent, the Raman score ranges from 0 to 1 based on the constituent’s weighted contribution to the tissue’s fingerprint spectra, but all scores sum to unity. High-wavenumber spectra were fit as a combination of spectral line shape functions^16^ and residual autofluorescence (**Supplement S1**). OH_score_ and CH_score_ represent the integrated areas of the OH and CH_2_ peaks, reflecting tissue hydration and organic components, respectively, normalized to sum to unity. After normalization, fingerprint scores possess two degrees of freedom, and high-wavenumber scores possess one degree of freedom.

For tissues with additional biochemical complexity, the multivariate regression model can incorporate additional terms to account for the spectral contribution of the added molecular constituent(s), such as a hydrogel scaffold in regenerating neocartilage or the contribution of subchondral bone in thinned cartilage samples.

### Tissue-mimetic phantom fabrication

Cartilage-specific biochemical phantoms were prepared by mixing chondroitin sulfate A sodium salt (bovine cartilage; C6737 Sigma) and gelatin (bovine skin, Sigma G9391) in deionized water. A total of 96 phantoms were produced spanning the following nominal GAG concentrations (0%, 0.9%, 1.8%, 2.7%, 3.6%, 4.5%, 5.5%, 6.4%, 7.3%, 8.2%, 9.1%, 10.0%) and gelatin concentrations (0%, 2.9%, 5.7%, 8.6%, 11.4%, 14.3%, 17.1%, 20.0%), with selected visually distinct phantoms shown in **Fig. 2C**. Component masses were recorded on a scale (Mettler Toledo ME204; ±0.1 mg) at each addition to achieve precise ground-truth composition measurements for model development (**Supplement S3**). After adding components, tubes were mixed on a tube rotator (30 rpm, 56°C, 2 hours) to yield a uniform solution (**Fig. 2A**). For spectral acquisition, a drop of each phantom was pipetted onto a stainless-steel substrate maintained at 4°C to prevent evaporation.

### Derivation of phantom calibration curve/transformation matrix

Fingerprint and high-wave Raman spectra were acquired for each phantom. Regression coefficients (scores) for constituents of the phantoms (GAG_score_, GEL_score_, OH_score_) were derived by reference-based multivariate regression analysis of the Raman spectra (**Supplement S3**). For each constituent, the relationship between constituent concentration and Raman scores was fit to a generalized second-order polynomial function with cross terms, yielding 10 regression coefficients (*β*_ij_ matrix):

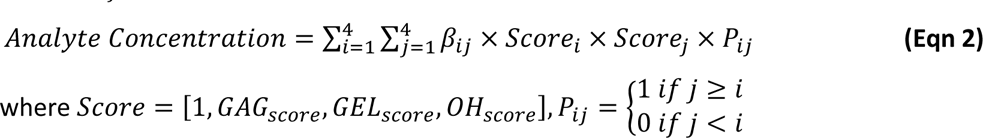

Negative concentration predictions were clipped to zero. Each constituent (GAG, Gelatin, water) has a unique and independent transformation curve. Determined parameters are listed in **Supplement S4**.

### Live cartilage explants

Cylindrical cartilage explants (∅3 mm) were aseptically cored from 2-month-old immature bovine femoral condyles (Green Village Packing Co). To minimize tissue heterogeneity’s effects on assay accuracy, the uppermost 2 mm of cartilage was excised, and cores were trimmed to 2 mm total thickness, yielding explants of exclusively deep zone/epiphyseal cartilage. Explants were cultured in basal medium consisting of high-glucose DMEM supplemented with 1 mM Na pyruvate, 50 µg/mL L-proline, 1% PS/AM antibiotic/antimycotic, 50 µg/mL ascorbate-2-phosphate, and 0.1% BSA. To induce osteoarthritis-associated composition changes, explants were cultured ±IL-1α at 50 ng/mL and ±DEX at 100 nM. Delayed IL-1α was administered for 7 days starting at day 14.

### Cartilage tissue assays

#### Raman

For explants not assayed in well plates, spectra were collected on both cylindrical faces for 30 seconds, processed to % wet weight as described above, and averaged.

#### Mechanical Properties

Explant thickness and diameter were measured with calipers, then the samples were compressed to 10% strain at 1μm/s between parallel impermeable platens while submerged in PBS, then allowed to relax for 900 seconds. Equilibrium modulus was calculated from the equilibrium load.

#### Biochemistry

Samples were weighed before and after lyophilization for water content determination. Lyophilized samples were digested in proteinase-K for 24 hours, and GAG content was quantified using a DMMB colorimetric assay. Protein comprises most non-GAG dry mass, of which type 2 collagen is the most important. Gelatin comprised predominantly of type 1 collagen is used as a spectral surrogate. Thus, generically collagen content (specified as COL*) was estimated indirectly as 100% - water% - GAG%.

#### Histology

Selected explants were fixed in 10% formalin, paraffin-embedded, sectioned (7 µm thick), and stained with hematoxylin, fast green, and safranin O^36^.

### Raman microscopy

Full-thickness bovine cartilage explants (∅5×2 mm) were harvested and treated with IL-1α to induce progressive proteoglycan degradation. Explants were collected after 0 (control), 5, 10, and 15 days (n = 4 per group). An additional hyaluronidase-treated group (3 mg/mL for 18 hours at pH 6.0 and 37**°**C) was included to generate severe GAG depletion (n = 3 per group). Samples were prepared by removing the subchondral bone, bisecting the cartilage sagittally, and imaging the exposed cross-section by Raman microscopy. Maps were generated with an integration time of 3 seconds per pixel. Adjacent transverse slices from the same explants were collected for matched depth-dependent DMMB GAG assay measurements, enabling Raman-derived GAG map comparisons with biochemical measurements from corresponding depth regions. Human samples underwent identical dissection for microscopy without culture.

### *In-vitro/ex-vivo* Raman monitoring of tissue explants

Bovine cartilage explants (∅5×2 mm) including the superficial zone were cultured in commercial 4-well plates equipped with Raman-transparent MgF_2_ windows adhered to the bottom of each well with silicone sealant (Dowsil 733) (**Fig.5B**). Explants were cultured for 10 days in control medium or treated with 10 ng/mL IL-1α with or without 100 ng/mL IGF-1, a disease-modifying osteoarthritis drug candidate (n = 3 per group). With the explant superficial surface in contact with the windows, Raman spectra were acquired daily through the MgF_2_ window with the plate lid closed. For comparison to conventional biochemical assays, parallel explants (∅3×2 mm) were cultured under identical conditions (n = 3 per group per time point) and collected at days 0, 4, 8, and 10 for endpoint Raman and biochemical analysis.

### *In vivo* Raman monitoring equine stifle joint

The arthroscopic Raman needle probe tip was sterilized by autoclaving. The probe body and optical fibers were covered with a sterile drape. Raman spectra were collected by inserting the needle probe through a parapatellar arthrotomy between the middle and lateral patellar ligaments. Follow-up Raman measurements and arthroscopic visual inspection were performed at 3 and 6 months post-repair. Twelve-month Raman measurements and macroscopic photographs were acquired postmortem on fresh tissue. Spectra were acquired at each site while holding the probe normal to the surface with minimal contact pressure. After baseline Raman acquisition, two 15 mm diameter chondral defects were created on the lateral trochlear facet at proximal and distal sites by removing all noncalcified cartilage to the calcified layer. Defects were immediately filled with cell-seeded collagen scaffolds^37^ and covered with a membrane held in place with resorbable sutures (**Fig. 6C**). Due to intermittent excessive autofluorescence in the high-wavenumber region, a simplified phantom calibration system was developed that used only fingerprint scores (**Supplement S7**).

### Statistical analysis

For **Fig. 6F**, significance was determined by two-way ANOVA of data categorized by time point and tissue type (repair versus surrounding native cartilage), followed by post hoc Tukey–Kramer testing (α = 0.05). Only differences between defect tissue and surrounding native cartilage are shown. Significant comparisons between other groups are not shown. Separate ANOVAs were performed for each of the two defects and two components (GAG and protein), yielding four tests in total.

## Acknowledgements

Research reported in this publication was supported by the National Institutes of Health (NIH) award number R01AR081393, the Arthritis Foundation, MTF Biologics, the Boston University Ignition Award, and Boston University Quantitative Biology & Physiology program (T32GM145455). The opinions, findings, and conclusions, or recommendations expressed are those of the authors and do not necessarily reflect the views of the National Institutes of Health.

