## Supplementary material for "Absolute Quantification of Tissue Composition by Raman Spectroscopy": Supplement.pdf

### Supplementary information

#### S1: High-wavenumber spectral analysis

After preprocessing, the spectrum is a vector of absolute intensities for each pixel in the high-wavenumber region, matched with a vector of wavenumbers for each pixel. Where $i$ is the index of a pixel in a spectrum, the preprocessed function is fit using least squares regression to the equation

$$Spectrum_i = Fluorescence_i + \sum_{i=1}^5 Peaks_i + Error_i$$

Fluorescence is a linear equation where for each pixel, the intensity ($Y$) of the fluorescence at that pixel is given by

$$Y = A \times Wavenumber + B$$

Where $A$ and $B$ are curve fit parameters to be found during regression.

There are five Raman peaks in the high wavenumber region detectable with our hardware, three ascribed to C-H bonds and two ascribed to O-H bonds. The intensity ($Y$) of a single peak at a given pixel is given by a Gaussian function

$$Y = h \times e^{-(\ln(2) \times x^2)}$$

Where $X$ is a coordinate transform of wavenumber space into a normalized Gaussian, given by:

$$X = \frac{Wavenumber - P_0}{w/2}$$

Where $h$, $P_0$, and $w$ are parameters estimated during regression. $h$ represents the height of the peak, $P_0$ represents the position of the center of the peak, and $w$ represents the full width at half maximum. There are five peaks, each with their own values of $h$, $P_0$, and $w$. During regression, $P_0$ and $w$ are constrained to a range based on the assumed identities of the peaks. The curve fitting constraints are given by the following table.

| Peak assumed location | assignment | Assumed full width at half maximum | Maximum deviation in $P_0$ from assumed location | Maximum deviation in $w$ from assumed full width at half maximum |
| --- | --- | --- | --- | --- |
| 2892 $\text{cm}^{-1}$ | C-H | 48 $\text{cm}^{-1}$ | 10 $\text{cm}^{-1}$ | 10 $\text{cm}^{-1}$ |
| 2942 $\text{cm}^{-1}$ | C-H | 43 $\text{cm}^{-1}$ | 10 $\text{cm}^{-1}$ | 10 $\text{cm}^{-1}$ |
| 2988 $\text{cm}^{-1}$ | C-H | 48 $\text{cm}^{-1}$ | 10 $\text{cm}^{-1}$ | 10 $\text{cm}^{-1}$ |
| 3205 $\text{cm}^{-1}$ | O-H | 103 $\text{cm}^{-1}$ | 70 $\text{cm}^{-1}$ | 320 $\text{cm}^{-1}$ |
| 3403 $\text{cm}^{-1}$ | O-H | 437 $\text{cm}^{-1}$ | 100 $\text{cm}^{-1}$ | 360 $\text{cm}^{-1}$ |

Values for O-H bonds were chosen based on a fit to pure water. Values for C-H bonds were chosen based on a fit of 20% gelatin in water. In practice the peaks rarely deviate from their assumed location and widths, the bounds are adjusted manually to make the regression algorithm robust against getting stuck in a local minimum. All five peaks and the fluorescence function are fit simultaneously, giving the regression 17 degrees of freedom. For each peak, its total value is given by the integral of the peak intensity function from $-\infty \text{ cm}^{-1}$ to $\infty \text{ cm}^{-1}$. The $\text{CH}_{\text{score}}$ is equal to the sum of the integrals of the C-H peaks divided by the sum of the integrals of all peaks. The $\text{OH}_{\text{score}}$ is equal to the sum of the integrals of the O-H peaks divided by the sum of the integrals of all peaks. This satisfies the normalization condition that $\text{CH}_{\text{score}} + \text{OH}_{\text{score}} = 1$, and removes the influence of the fluorescence fit from the final values. The individual peaks with the found $h$, $w$, and $P_0$ are shown in the spectrum graph with dashed lines.

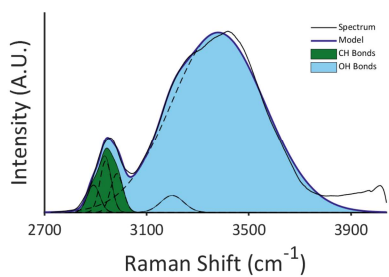

**Figure S1:** Highwave spectrum regression

### S2: Spectral comparison of gelatin and type-II collagen

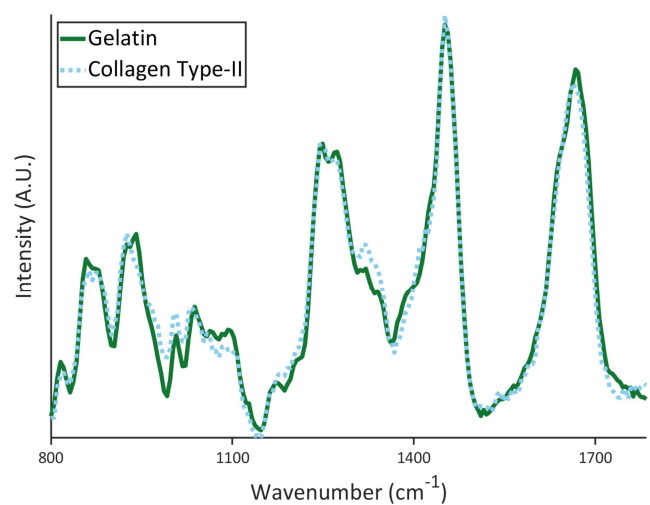

**Figure S2:** Comparison of preprocessed Raman fingerprint spectra of gelatin from bovine skin and collagen type II from chicken sternum.

#### **S3: Phantom composition data**

Phantom compositions are provided in the .CSV file in the supplement .ZIP folder

Values are expressed as mass fractions (g/g) from 0 to 1. For example, 90% water corresponds to a water mass fraction of 0.9.

##### S4: Raman calibration coefficients

The standard curve is described by the equation

$$\text{Analyte Concentration} = \sum_{i=1}^4 \sum_{j=1}^4 \beta_{ij} \times \text{Score}_i \times \text{Score}_j \times P_{ij}$$

##### (Eqn 2)

where

$$\text{Score} = [1, \text{GAG}_{\text{score}}, \text{Protein}_{\text{score}}, \text{OH}_{\text{score}}], P_{ij} = \begin{cases} 1 & \text{if } j \geq i \\ 0 & \text{if } j < i \end{cases}$$

The long form of this equation is represented as:

$$\begin{aligned} \text{Analyte Concentration (\% ww)} = & 100\% * (\beta_{1,1} * 1 * 1 + \beta_{1,2} * 1 * \text{GAG}_{\text{score}} + \beta_{1,3} * 1 * \text{Protein}_{\text{score}} + \beta_{1,4} * 1 * \text{OH}_{\text{score}} + \\ & \beta_{2,2} * \text{GAG}_{\text{score}} * \text{GAG}_{\text{score}} + \beta_{2,3} * \text{GAG}_{\text{score}} * \text{Protein}_{\text{score}} + \beta_{2,4} * \text{GAG}_{\text{score}} * \text{OH}_{\text{score}} + \beta_{3,3} * \text{Protein}_{\text{score}} * \text{Protein}_{\text{score}} + \\ & \beta_{3,4} * \text{Protein}_{\text{score}} * \text{OH}_{\text{score}} + \beta_{4,4} * \text{OH}_{\text{score}} * \text{OH}_{\text{score}}) \end{aligned}$$

The curve-fit tuning parameters are reported in the table below. Calculations use mass fractions in grams per gram rather than percentages. For example, 90% water is represented as 0.9 (grams of water per gram of phantom).

Reported values are multiplied by 100 to give percent wet weight.

| Beta matrix index | Associated term | Value for GAG measures | Value for Protein measures | Value for Water measures |
| --- | --- | --- | --- | --- |
| $\beta_{1,1}$ | $1 * 1$ | 3.952206 | 1.834712 | -4.63305 |
| $\beta_{1,2}$ | $1 * \text{GAG}_{\text{score}}$ | -0.49731 | 0.955563 | -0.51702 |
| $\beta_{1,3}$ | $1 * \text{Protein}_{\text{score}}$ | -0.54039 | 0.521648 | -0.0263 |
| $\beta_{1,4}$ | $1 * \text{OH}_{\text{score}}$ | -8.02585 | -4.31649 | 12.05618 |
| $\beta_{2,2}$ | $\text{GAG}_{\text{score}} * \text{GAG}_{\text{score}}$ | 0.21063 | 0.098384 | -0.30756 |
| $\beta_{2,3}$ | $\text{GAG}_{\text{score}} * \text{Protein}_{\text{score}}$ | 0.48723 | 0.590386 | -1.0723 |
| $\beta_{2,4}$ | $GAG_{score} * OH_{score}$ | 0.51741 | -1.0666 | 0.607842 |
| $\beta_{3,3}$ | $Protein_{score} * Protein_{score}$ | 0.037623 | 0.410085 | -0.44516 |
| $\beta_{3,4}$ | $Protein_{score} * OH_{score}$ | 0.528641 | -0.69595 | 0.210689 |
| $\beta_{4,4}$ | $OH_{score} * OH_{score}$ | 4.07889 | 2.514704 | -6.46148 |

### S5: Conventional assays applied to phantom solutions

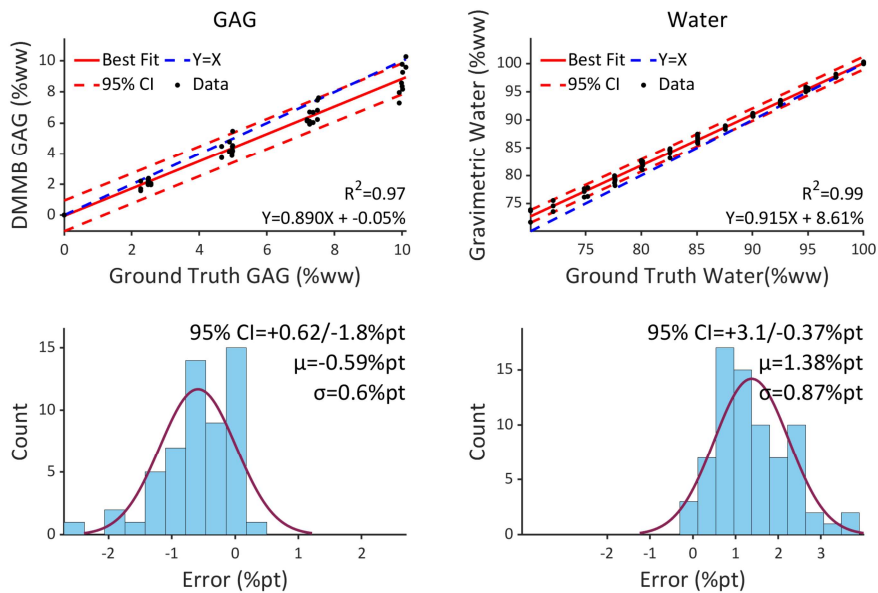

**Figure S5:** Comparison of phantom ground truth composition and biochemically assayed composition. The 95% confidence intervals expressed as percent of the sample weight do not match the uncertainty in the data. This is visible as the red dashed 95% confidence intervals calculated using the methods for phantoms don't match the observed spread in the data. The error in water is a multiple of the dry weight and the Error in GAG is a multiple of the GAG content. Assays were performed on a set of 25 phantoms and performed in triplicate, with outliers (single assays in a triplicate set with very different values) excluded. Assay performed by a scientist with 10 years of wet lab experience, correlations are typically worse for less experienced workers due to inconsistent technique and a higher incidence of mistakes.

### **S6: Video demonstration of real time analysis**

Captions for video file in supplement .zip folder:

The video is the of the surgical probe being used in a similar manner to actual surgery (held in the hands with overhead lights dimmed). The target is an immature bovine distal femur with a focal defect created by enzymatic GAG depletion using trypsin. The probe being used for single points, alternating back and forth between points in the defect and points in the healthy surrounding tissue. When the button on the probe is pressed or a foot pedal is used, a spectrum is collected at a point and analyzed with the results displayed on the UI in %pt.

### S7: Simplified fingerprint-only model for in vivo measurements

Due to extreme fluorescence in the surgical spectra making highwave analysis unreliable, a simplified phantom model was developed which does not use highwave data. The simplified model is derived from equation 2. Model performance is shown in Supplement S10.

$$\text{Analyte Concentration} = \sum_{i=1}^3 \sum_{j=1}^3 \beta_{ij} \times \text{Score}_i \times \text{Score}_j \times P_{ij}$$

Where

$$\text{Score} = [1, \text{GAG}_{\text{score}}, \text{Protein}_{\text{score}}], P_{ij} = \begin{cases} 1 & \text{if } j \geq i \\ 0 & \text{if } j < i \end{cases}$$

In long form this evaluates to:

$$\text{Concentration (\%weight)} = 100\% * (\beta_{1,1} * 1 * 1 + \beta_{1,2} * 1 * \text{GAG}_{\text{score}} + \beta_{1,3} * 1 * \text{Protein}_{\text{score}} + \beta_{2,2} * \text{GAG}_{\text{score}} * \text{GAG}_{\text{score}} + \beta_{2,3} * \text{GAG}_{\text{score}} * \text{Protein}_{\text{score}} + \beta_{3,3} * \text{Protein}_{\text{score}} * \text{Protein}_{\text{score}})$$

The fitted coefficients are:

| Beta matrix index | Associated term | Value for GAG measures | Value for Protein measures | Value for Water measures |
| --- | --- | --- | --- | --- |
| $\beta_{1,1}$ | $1 * 1$ | -0.00816 | -0.01474 | 1.022904 |
| $\beta_{1,2}$ | $1 * \text{GAG}_{\text{score}}$ | 0.252029 | 0.127977 | -0.38001 |
| $\beta_{1,3}$ | $1 * \text{Protein}_{\text{score}}$ | 0.034096 | 0.151554 | -0.18565 |
| $\beta_{2,2}$ | $\text{GAG}_{\text{score}} * \text{GAG}_{\text{score}}$ | -0.1376 | -0.20905 | 0.346647 |
| $\beta_{2,3}$ | $\text{GAG}_{\text{score}} * \text{Protein}_{\text{score}}$ | 0.096093 | 0.029882 | -0.12598 |
| $\beta_{3,3}$ | $\text{Protein}_{\text{score}} * \text{Protein}_{\text{score}}$ | -0.02548 | 0.048206 | -0.02272 |

### S8: Performance of the simplified fingerprint-only model

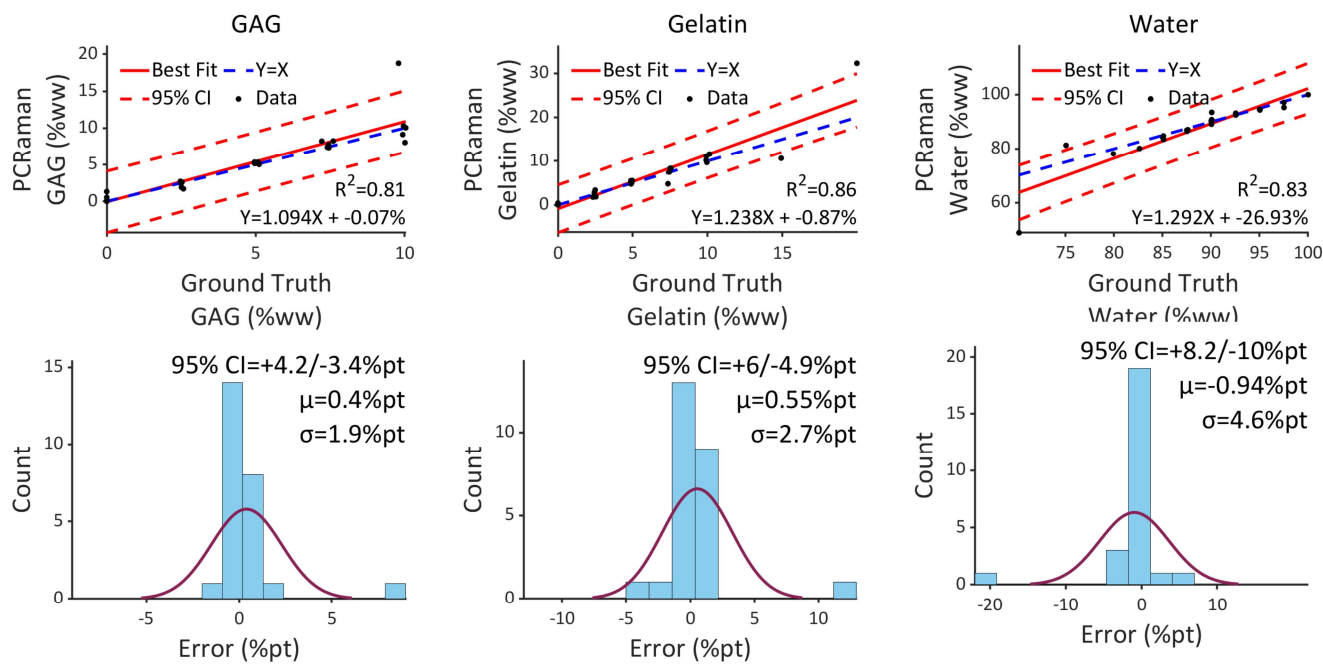

**Figure S8:** Phantom data for simplified model using only fingerprint scores to be used in surgery. Only 25 phantoms are used as the curve fit has fewer regression parameters to constrain. Certain key points around the boundary of composition space have an outsized effect on model accuracy. These points create outliers when left out in validation but are not left out for the final model used on tissue. Using only 25 points rather than 96 exaggerates the outlier, making the 95% CI appear much wider than it would be in practice where key points are not excluded. Excluding the single outlier point results in similar to but slightly worse performance than the full phantom model which includes highwave data.

#### S9: Comparison of representative spectra from an explant and a phantom

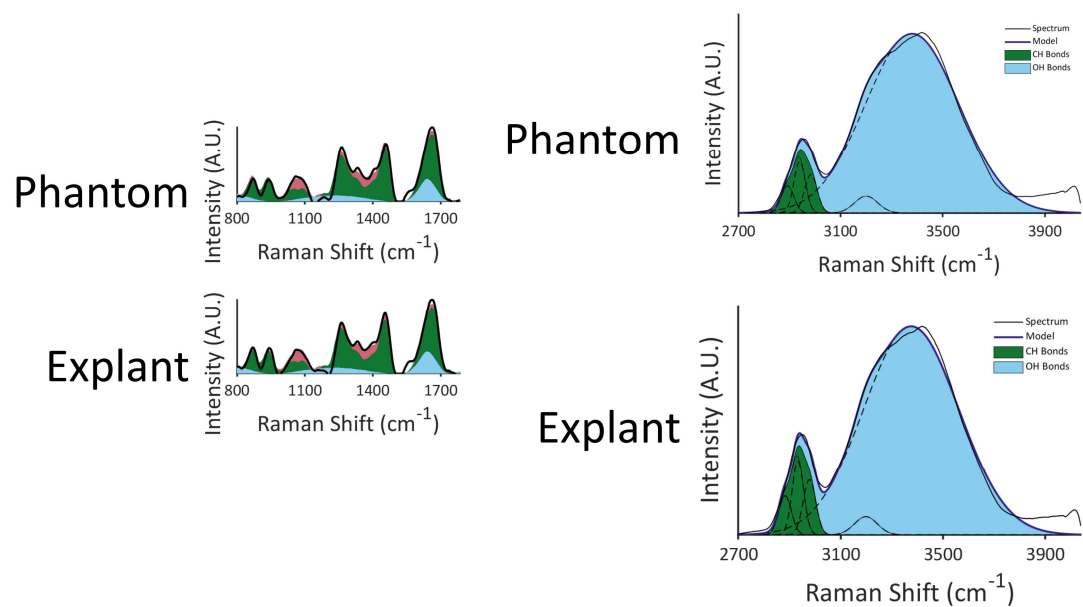

Figure S9: Representative spectra of a phantom and explant of similar composition showing similar spectra
